# Particle picking and centering in cryo-electron micrographs via a vanishing radial kernel

**DOI:** 10.64898/2026.11.02.763610

**Authors:** Ayelet Heimowitz, Nir Sharon

## Abstract

A training-free procedure combines particle picking and approximate centering in cryo-electron microscopy through a single convolution of the micrograph with a compact radial kernel, whose weights increase with distance from the origin, saturate in an annulus, and vanish beyond it. Particles are darker than their surroundings and much weaker than the noise; each carves a well in the response, and local minima of the response are candidate centers. Constant background offsets leave these extrema unchanged away from image boundaries, and a local stability bound controls the displacement of a well’s minimum under signed perturbations without assuming a symmetric particle. Grid seeds are iteratively updated, nearby endpoints are merged, and two-dimensional classification follows. On the *Plasmodium falciparum* 80S ribosome and *β*-galactosidase data sets, reconstructions reach 3.68 and 4.30 Å, compared with 3.77 and 4.36 Å using reference coordinates, with picking at 0.20–0.26 s per micrograph.

## 1 Introduction

Single-particle cryo-electron microscopy (cryo-EM) reconstructs the three-dimensional structure of a macromolecule from tens of thousands to millions of noisy two-dimensional projections (Frank, 1996). Identifying these projections in the micrographs, a task known as *particle picking*, remains one of the most labor-intensive steps of the cryo-EM computational pipeline.

Classical particle pickers correlate the micrograph with templates or with fixed functions such as the difference of Gaussians (Voss et al., 2009), and the statistics of such detectors are well understood (Sigworth, 2004). Template-based picking combined with sorting and 2D classification is available in standard software packages (Scheres, 2015). Other approaches train a binary classifier from examples selected automatically from the micrograph (Heimowitz et al., 2018), and widely used learning-based pickers rely on neural networks trained on annotated or partially labeled data (Bepler et al., 2019; Wagner et al., 2019). These approaches differ in their requirements for references, training data and computational resources. Even when a particle is detected correctly, the returned coordinates can leave it off-center in the extracted image.

Centered projections matter downstream. Once the picker has returned the coordinates of putative projections, projection images are extracted from the micrograph. These images are often normalized using pixels near their edges, which implicitly assumes the projections are approximately centered so the edges predominantly contain noise. Reference-free 2D classification and 3D refinement likewise search only a limited range of in-plane translations for each projection image (Scheres, 2012; Punjani et al., 2017; Zivanov et al., 2018; Kimanius et al., 2021), again implicitly assuming that the extracted projections are approximately centered. Poorly centered projections can enlarge the search, blur class averages when their offsets remain uncorrected, and be discarded as outliers; conversely, centering the images before reconstruction lets more projections contribute, which can improve reconstruction resolution (Heimowitz et al., 2021). Centering, however, is usually done after particle picking, separately for each projection image, which is costly with many projections.

In this paper, we propose a method that picks approximately centered projections: we perform picking and centering together, in a single step, directly on the micrograph. Each micrograph is filtered once with a non-negative, radially symmetric kernel that grows linearly with the distance from its origin up to the ramp radius *r*_0_ = *R/*2, remains constant up to *L* = 7*R/*8, and then vanishes. In the recorded micrograph, particle interiors are predominantly darker than their surroundings, and the kernel is non-negative. Thus, each particle carves a well in the response, providing a centering landscape in which local minima serve as candidate particle centers. This construction is motivated by the surrogate center-of-mass landscape in (Heimowitz et al., 2021). Specifically, the noise enters the response additively, and the kernel filters it. We bound its effect on a response minimum when the filtered perturbation has sufficiently small local oscillation. Our analysis includes local centering properties and appears in §3. Because the method computes the response for the whole micrograph via a single FFT-based convolution and requires no training or templates, it is fast, taking a fraction of a second per micrograph on the reported hardware.

Our main contribution is embedding a centering criterion into a training-free, micrograph-wide procedure rather than centering each extracted projection separately; here, a single micrograph correlation yields a landscape used to locate approximately centered particle candidates through-out the micrograph. Specifically, we mark the following:

1. **A centering landscape for the whole micrograph**. We introduce the vanishing radial kernel and evaluate the resulting local centering landscape at every pixel via a single FFT-based convolution, producing a response map whose relationship to the surrogate center-of-mass landscape of (Heimowitz et al., 2021) we discuss in §3.
2. **Analysis of the response map**. We establish a quantitative local stability bound for response minima under controlled signed perturbations, without assuming a symmetric particle. We also discuss the effects of signed profiles and of the background, including exact invariance to constant noise offsets away from image boundaries and the role of the particle size relative to the kernel. More in §3.
3. **A training-free picker that returns approximately centered coordinates**. A regular grid of seed positions initializes minimization of the response,^1^ and coincident endpoints are merged (§2.3). The scale *R* is the only user-selected spatial parameter, the response-map computation does not depend on the number of particles, and the same descent can re-center coordinates produced by any other picker. The procedure deliberately favors recall and leaves false positives to routine 2D classification, as is common practice in cryo-EM (Scheres, 2015; Zivanov et al., 2018).
4. **Validation on experimental data**. On EMPIAR-10028 and EMPIAR-10017, reconstructions from our picks reach resolutions comparable to those obtained from the deposited coordinates, with picking times of 0.2–0.26 s per micrograph; on EMPIAR-10420, a nearest-neighbor coordinate comparison assesses agreement with the polished particles retained in the original study (see §4).

## 2 Methods

### 2.1 Image formation model and contrast convention

For a micrograph *I* on an *N* × *N* pixel grid, consider the model

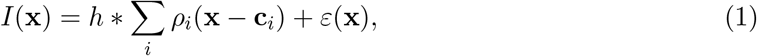

where *ρ*_*i*_ is a particle projection, **c**_*i*_ its position, *h* the point-spread function, and *ε* noise (Frank, 1996). In recorded micrographs, the CTF at the defocus values used renders protein darker than its surroundings. That is, the noise term which is not affected by the CTF is larger in magnitude than the particle term.

The micrograph is used as recorded, and no contrast inversion is applied. A spatially constant offset in *ε* adds only a constant to the response away from image boundaries and therefore leaves its extrema unchanged. Spatial fluctuations can alter the response minima. In §3, we bound their effect under a local oscillation condition. CTF phase flipping may precede the procedure but is not required; it does not make the underlying particle profile non-positive, so the non-positive profile used in §3 is an idealization.

We use *R* as the prescribed spatial scale for grid seeding, neighborhood searches, and endpoint merging. It is chosen to be comparable to the particle radius in the working image (§2.2). The kernel ramp and support radii are derived from *R* in §2.2. We denote the closed Euclidean disc of radius *r* by *B*_*r*_(**x**). Coordinates and radii are measured in pixels unless stated otherwise. Integrals describe the continuous response; implementation uses pixel sums.

### 2.2 Kernel and response map

Set *r*_0_ = *R/*2 and *L* = 7*R/*8 = 1.75*r*_0_, and define

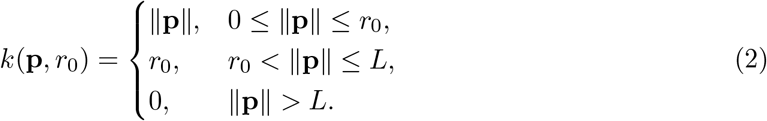

The kernel is a distance ramp, a constant annulus, and a sharp outer cutoff (Fig. 1). The response is

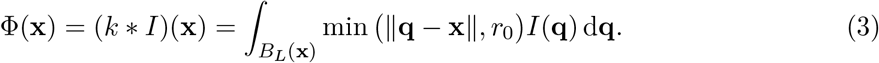

**Figure 1:**
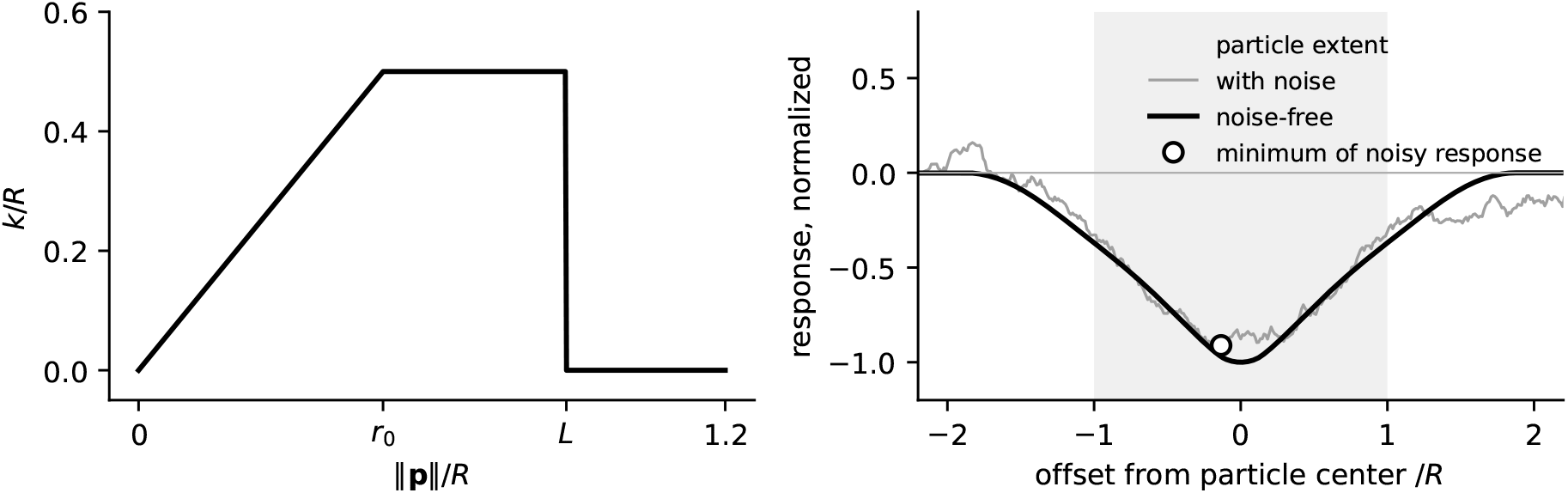
Left: radial kernel profile, with distances and weights divided by *R*. Right: response along a line through the center of a particle that is darker than its surroundings, computed on a pixel grid with *R* = 82 for the projection of a uniform ball of radius *R* (shaded extent). Black: noise-free. Gray: the same micrograph with white noise ten times larger than the particle contrast. The circle marks the minimum of the noisy response along this line, 11 pixels (0.13*R*) from the true center. The response is shown relative to its background level and normalized by the well depth. The particle carves a well whose minimum is at its center, and the well persists under the noise. The curves do not represent a CTF correction or a detection guarantee.

Because *k* is symmetric, convolution and correlation coincide. In a discrete implementation, the kernel origin is placed at zero lag, and padding or the exclusion of border pixels prevents wraparound from affecting retained coordinates.^2^

Because *k* is non-negative and the particle term is smaller than the ice term, the particle lowers the response wherever *B*_*L*_(**x**) overlaps it: its contribution to Φ is a well of extent about *L* plus the particle radius, and it vanishes beyond, since the outer cutoff *L* makes the response local. The noise contributes *k* * *ε* to the response. Its constant part shifts Φ uniformly, whereas the filtered fluctuations can change the locations, depths, and existence of the particle wells. Where in the well the minimum lies depends on the particle size relative to the kernel. The scale *R* is chosen so that *L* is comparable to the particle radius. The annulus, which carries the largest weight *r*_0_, then lies over the particle when the kernel is centered on it, and moving the kernel exchanges annulus pixels over the particle for background pixels, which raises the response; the well is deepest at the center. If instead the whole particle lay inside the ramp 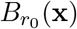, it would receive the smallest weights, and the response would be locally maximal, not minimal, at its center (§3); a kernel much larger than the particle is therefore excluded by the choice of *R*. Under sufficiently small signed perturbations, a nearby minimum persists (Proposition 1). Response values alone do not distinguish a particle well from a well caused by noise, contamination or thicker ice, and particle centers need not be global minima.

Figure 2 illustrates the above. We show a clean and noisy hedgehog silhouette, darker than the background as in a micrograph. In each of these, we mark in blue the reference center, in yellow a small shift from the center, and in pink a large shift. Additionally, we present the kernel. When the kernel is centered on the hedgehog, its annulus lies over silhouette pixels, which receive the largest weight *r*_0_ and lower the response the most. A small shift moves part of the annulus onto background pixels and raises the response. If *B*_*L*_(**x**) contains no hedgehog pixels, the particle contribution to the response is zero. The end result (in the noiseless case) can take the form shown in Figure 1.

**Figure 2:**
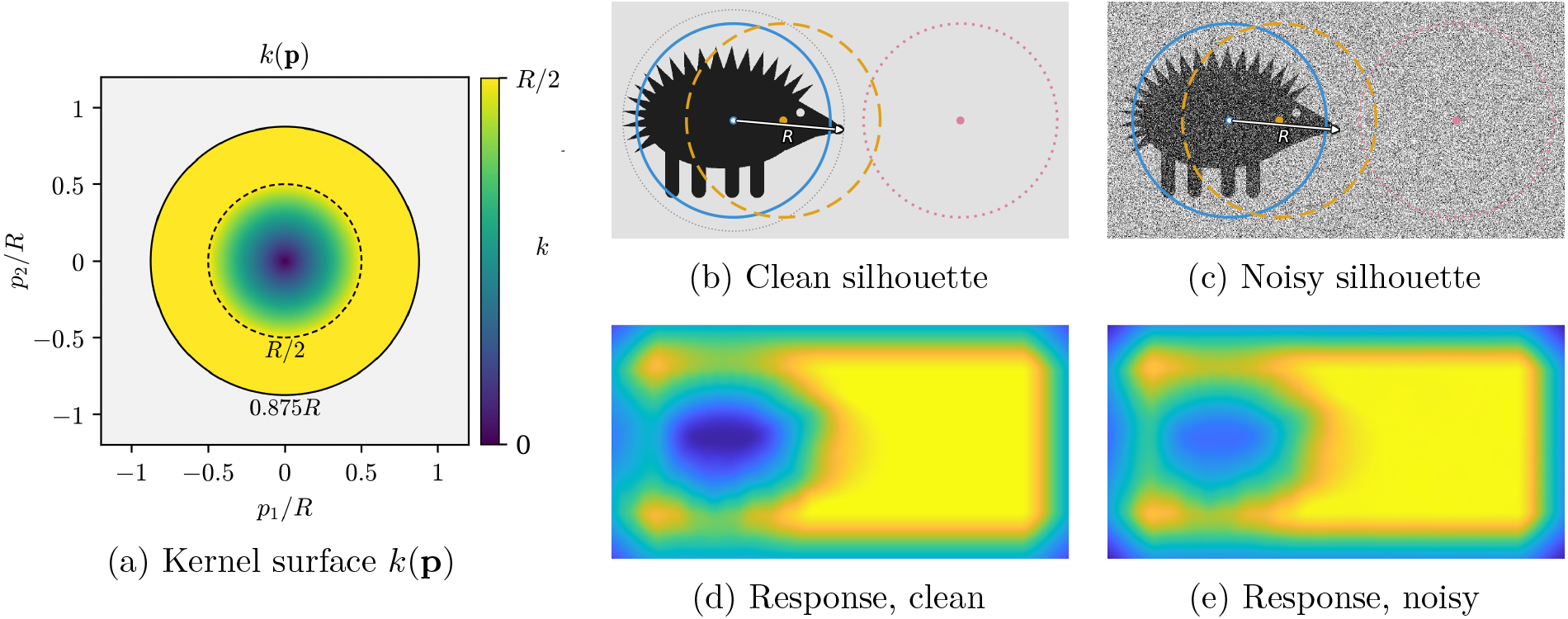
Radial kernel and its response on clean and noisy hedgehog silhouettes. a: the kernel *k*, which grows linearly from the origin up to *r*_0_ = *R/*2, stays constant up to *L*, and vanishes beyond (see also Figure 1). b, c: silhouettes with three disks of radius 0.875*R*, centered at the center of mass (solid), slightly shifted (dashed), and far away so that the disk contains no hedgehog pixels (dotted). d, e: the corresponding responses, drawn directly below each silhouette at the same scale.

### 2.3 Grid-seeded picking and centering

Picking proceeds by searching for local minima of Φ from grid seeds (Algorithm 1). The pseudocode uses the micrograph as recorded, with particles darker than the background, so that particles correspond to wells of Φ. The micrograph is covered by a regular square grid of seed positions with spacing 2*R*. Each seed is treated as an initial estimate of a projection center. The seed is then iteratively moved to the pixel with the lowest response in the *R* × *R* window *W*_*R*_(**x**) centered at its current position **x**, restricted to valid image pixels. An update is accepted only if it strictly decreases the response. Iteration stops when the current pixel itself attains the minimum in its search window. On a finite image grid, strict decrease ensures termination. Because many seeds reach the same or nearby endpoints, endpoints closer than a merge distance of 2*R* are merged to reduce duplicate picks.

Each seed stops at a minimum of Φ over its search window. Which particle centers are recovered depends on both the seed positions and the neighborhood updates. The procedure also returns response minima caused by noise, contamination, or carbon, i.e., false positives. This trade-off is deliberate: we prefer recall and use reference-free 2D classification to filter false positives, as is common practice in cryo-EM (Scheres, 2015; Kimanius et al., 2021). Figure 3 summarizes the response landscape and the resulting particle picking.

**Figure 3:**
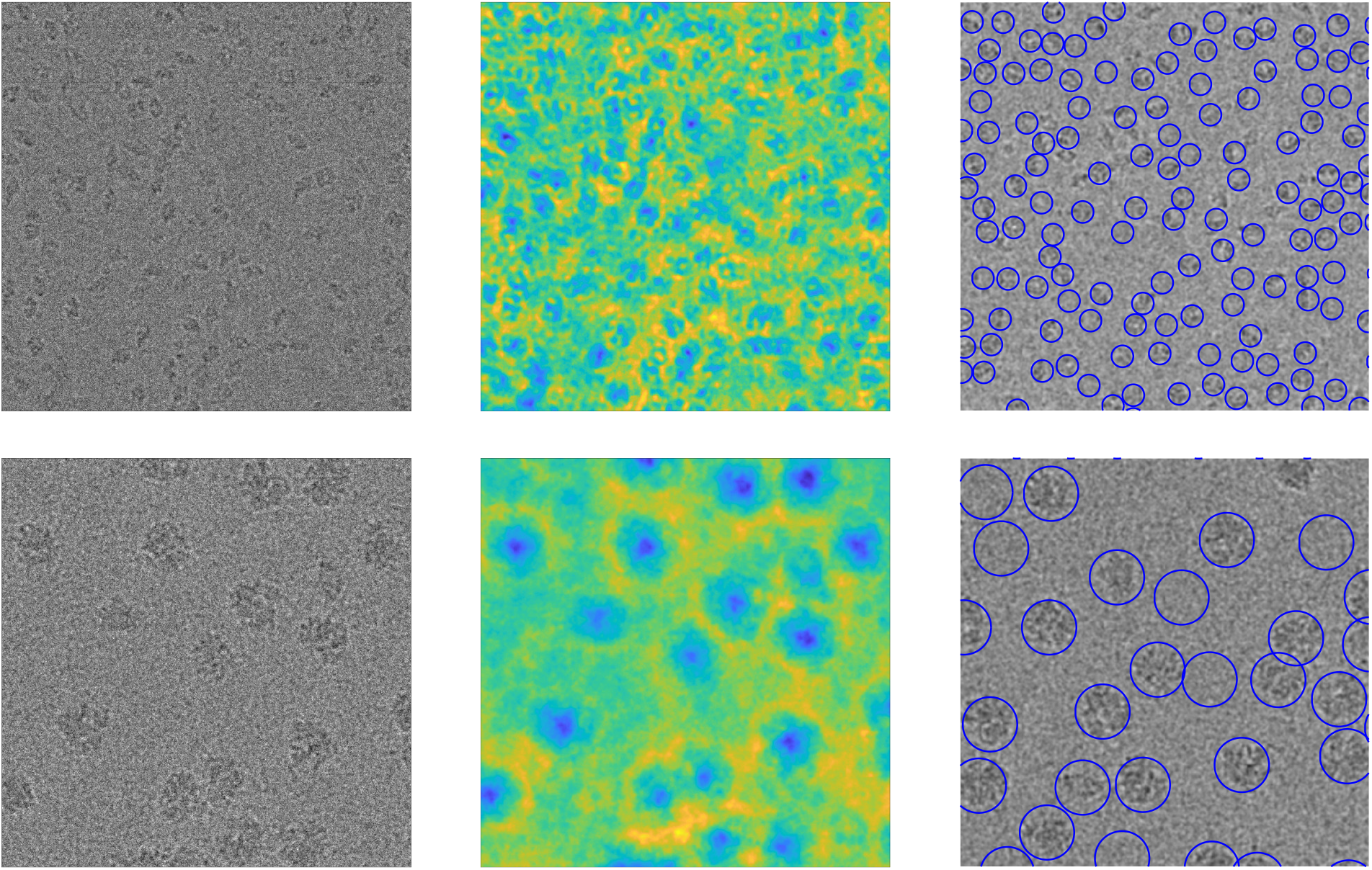
Examples of the proposed framework on EMPIAR-10017 (top row) and EMPIAR-10028 (bottom row). Left: cropped experimental micrographs without phase flipping. Middle: corresponding response maps. Right: selected particle coordinates.

#### Algorithm 1

Grid-seeded picking and centering.

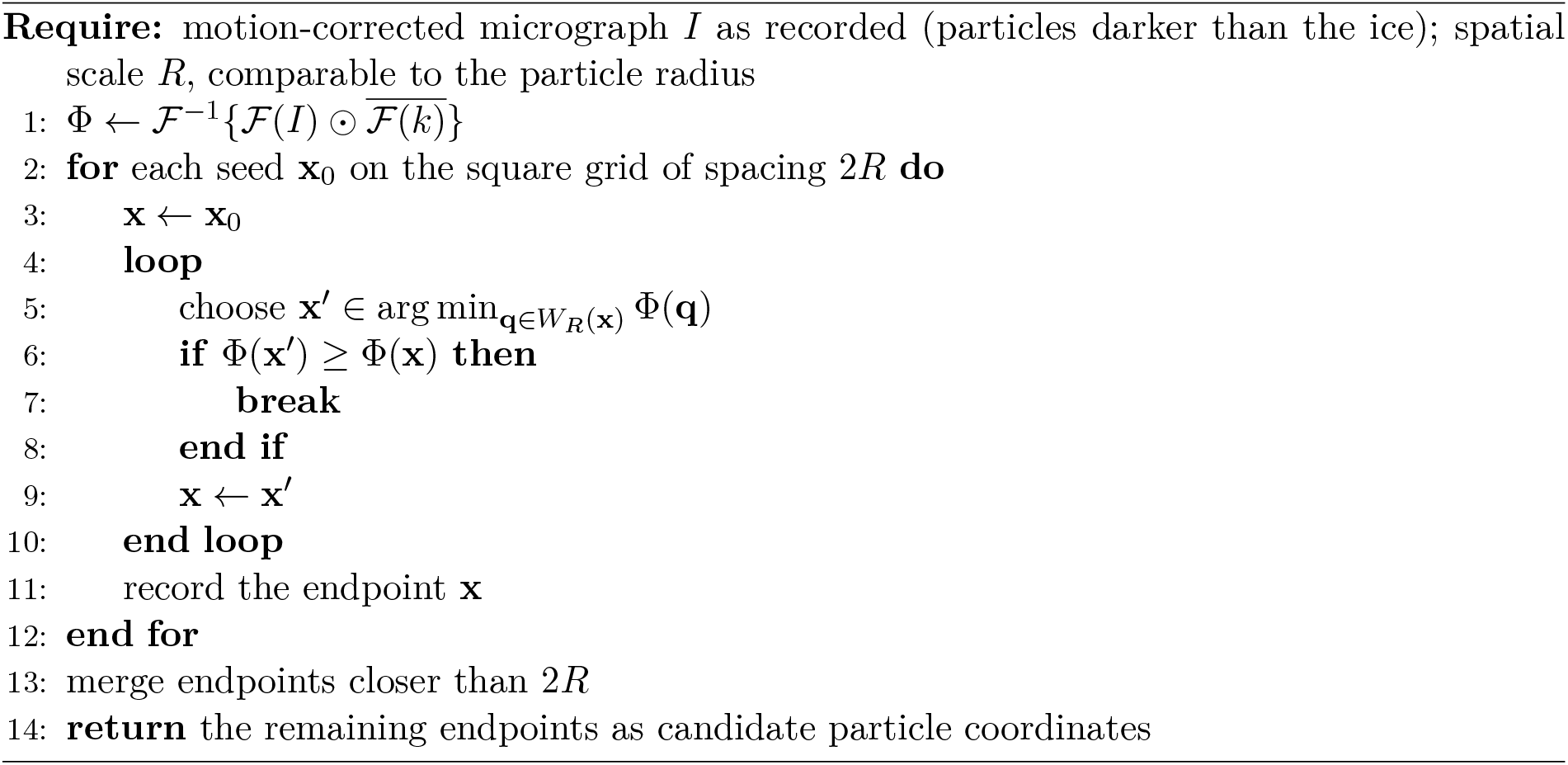

#### Centering

To center particles already picked by another method, we replace the grid seeds with the existing particle coordinates. The remainder of the method is unchanged.

## 3 Analysis of the centering landscape

### 3.1 Well geometry, the geometric median, and stability

Consider one particle of non-negative projected density *m* with nonzero mass, as it appears in *I* after convolution with *h*. Its dark contrast means that it enters the micrograph as −*m*, so its noise-free response is

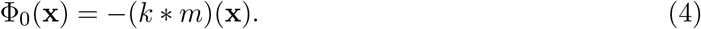

The particle is positive; the minus sign is its contrast. Minimizing Φ_0_ places the kernel where it covers the most of *m*, weighted by *k*. Since the weights are smallest near the kernel origin and equal to *r*_0_ in the annulus, the minimum favors positions at which the annulus lies over the particle.

The geometric median **g** of *m* minimizes

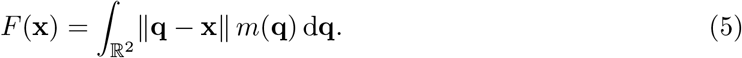

If supp *m* lies strictly inside 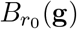, then (*k* * *m*)(**x**) = − *F* (**x**) and Φ_0_(**x**) = *F* (**x**) for all **x** near **g**, so **g** is a local *maximum* of Φ_0_, and minimization moves the particle into the annulus. A kernel whose ramp contains the whole particle therefore does not center it. This is why *R* is matched to the particle, so that the ramp lies inside the particle and *L* is comparable to its radius; in that regime the well is deepest at the center (Figure 1), as the curvature formula of §3.2 makes precise for radial profiles.

The relation to the surrogate center-of-mass landscape of (Heimowitz et al., 2021) is local. Integrating the enclosed mass of *m* over radii up to *r*_0_ gives the cone term *C*(**x**) = ∫ (*r*_0_ − ∥**q** − **x**∥)_+_*m*(**q**) d**q**. If *B*_*L*_(**x**) contains the whole particle, then (*k* * *m*)(**x**) = *r*_0_∫ *m* − *C*(**x**), so Φ_0_ and *C* differ by a constant. In the operating regime, however, the particle is not contained in *B*_*L*_(**x**), the mass inside *B*_*L*_(**x**) varies with **x**, and noise and neighboring particles add further terms; the two landscapes need not agree. Guarantees for the earlier surrogate therefore do not transfer automatically.

The next result quantifies the stability of the minimum of a well under a signed perturbation. It allows an arbitrary particle shape, requires no symmetry, and concerns a minimum of the landscape, not convergence of Algorithm 1.

#### Proposition 1

(Local centering under perturbations). *Let* Φ_0_ *be a continuous function with a local minimizer* **c**, *and suppose that for some s >* 0 *and λ >* 0

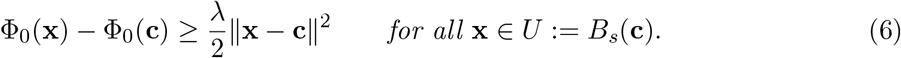

*Suppose that, on U, the observed response is* Φ = Φ_0_ + *E, where E is continuous, and put*

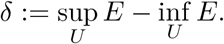

*If δ < λs*^2^*/*2, *every minimizer* **ĉ** *of* Φ *on U satisfies*

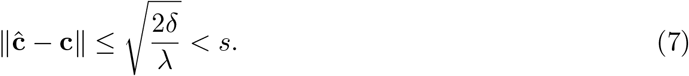

*Consequently, these minimizers lie in the interior of U, and* Φ *retains a local minimum near* **c**.

The proof is given in Appendix A.

Here Φ_0_ is the noise-free response of the particle, and *E* collects the noise response *k* * *ε* together with residual background, CTF mismatch, or neighboring signal. A constant component of *E* does not contribute to *δ*. Spatially varying perturbations can shift or eliminate the well, and Proposition 1 applies when their oscillation satisfies *δ < λs*^2^*/*2. A bound ∥*E* ∥_*∞,U*_ ≤ *ϵ* implies *δ*≤ 2*ϵ*. The growth condition (6) holds whenever Φ_0_ is twice continuously differentiable near **c** with ∇^2^Φ_0_(**c**) ⪰ *λ*_0_**I** for some *λ*_0_ *>* 0: for every *λ < λ*_0_ there is an *s >* 0 for which (6) holds. For a radial profile, Remark 1 gives *λ*_0_ explicitly. The estimate is conservative, weakens as the well bottom flattens, and does not assert uniqueness of the perturbed minimum. For an asymmetric particle, the minimizer **c** of the well need not coincide with the center of mass of the projection.

### 3.2 Signed profiles and background

Radial symmetry guarantees stationarity at the particle center. For signed or extended profiles, the curvature also depends on the profile near the kernel boundaries.

*Remark* 1 (Curvature for a signed radial profile). Let 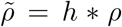 be the twice continuously differentiable image of a particle, radially symmetric about **c**, and write 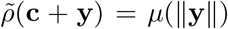, where *µ* may take either sign; in the idealization used here a particle has *µ* ≤ 0, while *ρ* itself is non-negative. Let 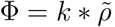, with *r*_0_ = *R/*2 and *L* = 7*R/*8. Then ∇Φ(**c**) = 0 and

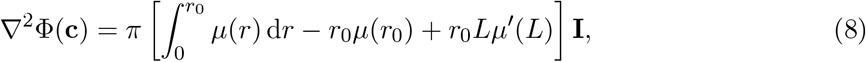

where *µ*′ denotes differentiation with respect to radius and **I** is the 2 × 2 identity matrix. A positive bracketed term gives a strict local minimum, and a negative term gives a strict local maximum. For a nonzero particle profile (*µ* ≤ 0) supported strictly inside 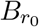(**c**), the boundary terms vanish, the bracket is negative, and the center is a strict local maximum, in agreement with §3.1. For a profile that extends beyond *L* and rises towards zero there, *µ*′(*L*) *>* 0, the cutoff term *r*_0_*Lµ*′(*L*) is positive; it outweighs the first two terms when the particle edge lies near *L*, and the center is then a strict local minimum. This is the operating regime, and *π* times the bracket is the constant *λ*_0_ of Proposition 1. A profile that is flat across *B*_*L*_(**c**) gives zero curvature and a flat well bottom, and CTF-induced oscillations can change the sign. The derivation is given in Appendix A.

The response is local: signal outside *B*_*L*_(**x**) does not affect Φ(**x**). Away from image boundaries, a constant offset *ε*_0_ adds only the constant *ε*_0_*K*, where *K* = ∫*k*, and leaves the extrema unchanged. Spatially varying noise can alter the minima. Proposition 1 controls its local oscillation after filtering. For example, an affine background *b*_0_ + ***β*** · **x** contributes *K*(*b*_0_ + ***β*** · **x**). A background gradient can therefore shift the response extrema.

### 3.3 What the bound says at a realistic noise level

To see how Proposition 1 and Remark 1 combine in practice, consider white noise with standard deviation *σ* per pixel and a particle of radius *R* whose image has central contrast *a*_*p*_ (the particle of Figure 1), so that *a*_*p*_*/σ* is the amplitude signal-to-noise ratio of a single pixel, well below one in cryo-EM. Three facts, derived in Appendix B, summarize the situation.

First, the mean of the noise, however large, adds only a constant to the response and does not move any extremum. Second, the kernel averages the noise fluctuations over about 2.4*R*^2^ pixels while it accumulates the particle coherently: the depth of the particle well exceeds the standard deviation of the noise response by a factor of about 1.1 (*a*_*p*_*/σ*)*R*, which is close to 9 for *R* = 82 pixels at *a*_*p*_*/σ* = 0.1. The well is therefore prominent in the response although the particle is invisible pixel by pixel; this is what the response maps of Figure 3 show. Third, when the filtered noise varies sufficiently little around the well, Proposition 1 bounds the displacement of its minimum. For *R* = 82 and *a*_*p*_*/σ* = 1*/*4, the illustrative bound is about 40 pixels, roughly half the particle radius. This estimate uses the explicit local noise condition stated in Appendix B.

In short, the large noise of recorded micrographs costs nothing in the location of the wells; it enters only through its averaged fluctuations, which Proposition 1 converts into a centering error that shrinks as the particle grows in pixels and as the signal-to-noise ratio improves. The bound treats the noise as adversarial near the center and is therefore conservative; actual displacements are expected to be smaller. Noise concentrated at low spatial frequencies is averaged less and requires a higher signal-to-noise ratio to achieve the same guarantee.

## 4 Results

### 4.1 Implementation and reconstruction workflow

We implemented the procedure in MATLAB and Python. The code is available on a designated repository at https://github.com/ayeltg/TRAP. The reported computing system comprised an Intel Core Ultra 7 CPU with 16 cores and an NVIDIA RTX 2000 GPU. Micrographs were used as deposited. Grid spacing and endpoint suppression distance were both *R*, and the kernel transform was computed once per dataset. We used RELION (Scheres, 2012; Zivanov et al., 2018; Kimanius et al., 2021) for particle extraction, classification, and refinement. We report resolutions at Fourier shell correlation (FSC) 0.143 between independently refined half-maps (Rosenthal and Henderson, 2003; Scheres and Chen, 2012).

The comparison is between complete candidate-selection workflows: our candidates received an additional round of 2D classification relative to the reference coordinates. The reported resolution differences therefore combine picking, selection, and centering effects. Table 1 reports the initial candidate counts, resolutions, and picking times.

**Table 1:** Picking and reconstruction results. Time is the reported mean picking time per micrograph, rather than total processing time; it excludes downstream classification and reconstruction. A dash indicates that no reference picking time is reported. The candidate-selection workflows differ in the number of 2D classification rounds.

| EMPIAR | Coordinates | Initial candidates | Resolution (Å) | Time (s) |
|---|---|---|---|---|
| 10028 | Reference | 158,212 | 3.77 | – |
|  | Our method | 252,920 | 3.68 | 0.20 |
| 10017 | Manual | 40,863 | 4.36 | – |
|  | Our method | 78,312 | 4.30 | 0.26 |

### 4.2 P. falciparum 80S ribosome (EMPIAR-10028)

EMPIAR-10028 contains 1081 micrographs of the *Plasmodium falciparum* 80S ribosome bound to emetine (Wong et al., 2014; Iudin et al., 2023). The original study selected 158,212 particles and retained 105,247 after classification; the latter are supplied as processed particles in 360 × 360-pixel boxes at 1.34 Å per pixel. These processed particles are distinct from the initial coordinate set used as the reference in Table 1.

The reference coordinates were processed using 25 iterations of 2D classification, initial-model building and 25 iterations of 3D classification, followed by re-extraction, refinement and post-processing. Our method, with *R* = 82 pixels, returned 252,920 candidates. Sixteen iterations of 2D classification retained 135,908, followed by a second 2D classification and the subsequent reconstruction steps. The resulting resolutions were 3.68 Å for our candidates and 3.77 Å for the reference workflow. The 3.2 Å resolution reported in (Wong et al., 2014) belongs to the original processing and is not the reference outcome reproduced here.

### 4.3 β-Galactosidase (EMPIAR-10017)

EMPIAR-10017 comprises 84 Falcon II micrographs of *β*-galactosidase (4096 × 4096 pixels, 1.77 Å per pixel), with manually selected coordinates (Scheres, 2015; Iudin et al., 2023). The particles are densely packed, making neighboring signal and endpoint suppression relevant. With *R* = 40 pixels, our method returned 78,312 candidates, approximately 932 per micrograph.

As for the ribosome, our candidates underwent two rounds of 2D classification, compared with one for the reference. The final particle counts were 32,993 and 32,240, respectively, with resolutions of 4.30 and 4.36 Å. Thus a larger initial candidate set produced a reconstruction of comparable resolution after classification; this is not a direct measurement of coordinate precision or recall.

### 4.4 Arabinosyltransferase EmbB (EMPIAR-10420)

A further coordinate comparison used the Falcon III subset of EMPIAR-10420, containing 2158 micrographs of arabinosyltransferase EmbB from *Mycobacterium smegmatis*, recorded at 0.665 Å per pixel (Tan et al., 2020). In the original study, lenient Gautomatch picking and RELION classification produced 162,271 candidates; further processing in cryoSPARC reduced this set to 19,755 before subsequent refinement and polishing (Tan et al., 2020; Punjani et al., 2017).

Figure 4 shows, for each deposited polished particle in the comparison, the distance to the nearest candidate from our picker. The histogram is concentrated at short distances relative to the 40-pixel marker. This one-sided nearest-neighbor comparison indicates proximity to refined reference coordinates, namely, to an estimated center-of-mass. Note that this comparison does not enforce one-to-one matches, quantify false positives or establish a direct centering accuracy.

**Figure 4:**
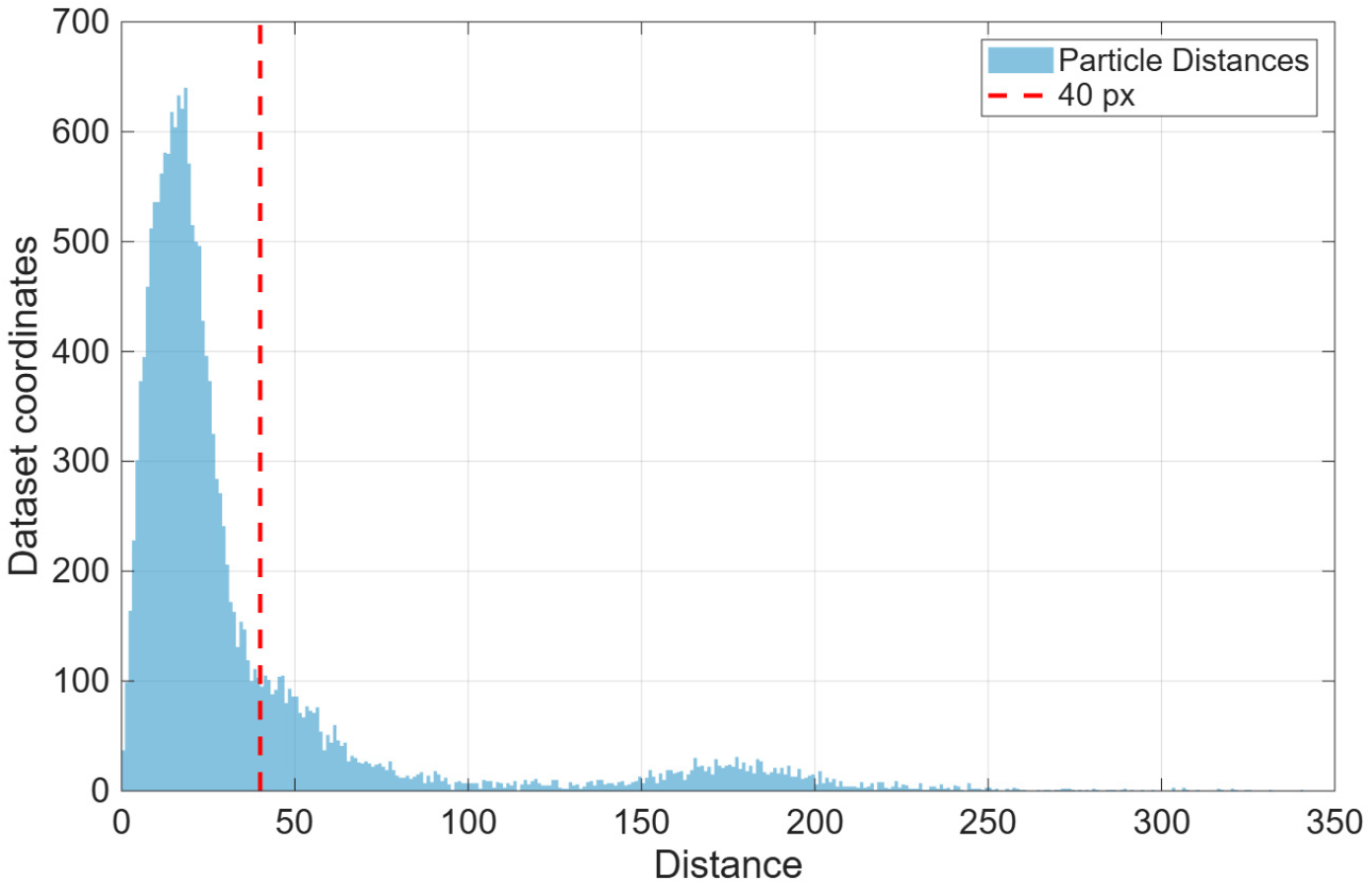
EMPIAR-10420: distances, in pixels, from deposited polished-particle coordinates to their nearest candidates from our method. The dashed line marks 40 pixels. This is a one-sided coordinate-agreement plot, not a precision–recall curve.

## 5 Discussion and conclusions

The radial response lets candidate picking and centering share a single micrograph-wide computation. Because the kernel is non-negative and particles are darker than their surroundings, each particle carves a well in the response; the noise, although much larger in magnitude, shifts the response by a constant and adds a smoothed perturbation whose effect Proposition 1 quantifies (§3.3). The geometric-median identity of §3.1 shows that the linear part of the kernel must lie inside the particle: a particle contained in the ramp is a local maximum, not a minimum, of the response. Proposition 1 bounds the displacement of a well’s minimum under controlled signed perturbations without assuming a symmetric particle, and (8) identifies the outer-cutoff term as the source of the central minimum for radial profiles.

Crowding, asymmetric mass, and CTF-induced tails can shift the response extrema; suppression at distance 2*R* can then discard neighboring particles. The scale *R* should therefore match the effective particle footprint in the working image: a cutoff *L* well inside the particle flattens the bottom of the well, a kernel that contains the particle inverts the central curvature, and a larger *R* increases neighboring contributions.

The experimental comparisons show that the candidate sets can support reconstructions with resolution comparable to the reference workflows, with reported picking times below 0.3 s per micrograph. The surplus candidates increase downstream classification cost, which is not included in those times. Matched classification protocols and quantitative coordinate matching would be needed to isolate centering improvements and establish detection recall and precision.

## Acknowledgements

N.S. is partially supported by NSF-BSF under grant 2024791, the BSF award 2024266, and the DFG award 514588180.

## Use of generative AI

Claude (Anthropic) assisted with optimizing the authors’ code and translating it from MATLAB to Python. Large language models, including ChatGPT (OpenAI), also assisted with manuscript organization and editing. The authors retain full responsibility for the manuscript, the code, and the validation and interpretation of the reported results.

## Data availability

The source micrographs and reference data are publicly available from the Electron Microscopy Public Image Archive under accession codes EMPIAR-10028, EMPIAR-10017 and EMPIAR-10420 (Iudin et al., 2023).

## A Mathematical proofs

*Proof of Proposition 1*. Compactness of *U* and continuity of Φ give a minimizer **ĉ** of Φ on *U*. Comparing with **c** and using Φ = Φ_0_ + *E*,

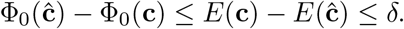

Combined with (6), this gives *λ* ∥**ĉ** − **c**∥^2^*/*2 ≤ *δ*, which is (7). Since 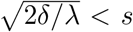, **ĉ** lies in the interior of *U* and is therefore a local minimum of Φ. If Φ_0_ is twice continuously differentiable near **c** with ∇^2^Φ_0_(**c**) ⪰ *λ*_0_**I**, then for any *λ < λ*_0_ continuity of the Hessian provides an *s >* 0 on which ∇^2^Φ_0_ ⪰ *λ***I**, and (6) follows from Taylor’s theorem with ∇Φ_0_(**c**) = 0.

**Derivation of** (8). Radial symmetry gives stationarity and a scalar Hessian. Differentiating the smooth profile under the convolution and integrating by parts on [0, *r*_0_] and [*r*_0_, *L*] gives

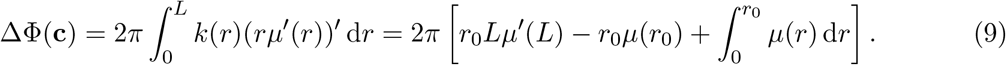

Half of this trace is each Hessian eigenvalue.

## B Calculations for the white-noise illustration

Throughout, *r*_0_ = *R/*2, *L* = 7*R/*8, lengths are measured in pixels and pixel area is one. Integrals approximate sums over the sampled kernel. Write *ε* = *ε*_0_ + *η*, where *ε*_0_ is constant, and *η* is spatially independent, zero-mean noise with variance *σ*^2^ per pixel. The constant contributes *ε*_0_*K* to the response away from image boundaries. The kernel satisfies

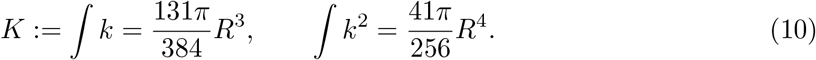

Consequently, the filtered fluctuation *E* = *k* * *η* has pointwise standard deviation

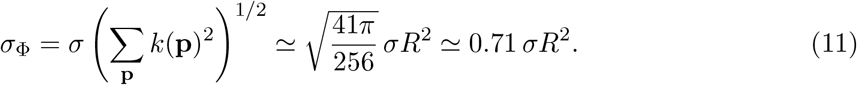

This variance calculation requires no Gaussian assumption. The filtered values at different locations are generally correlated. For the idealized particle, set

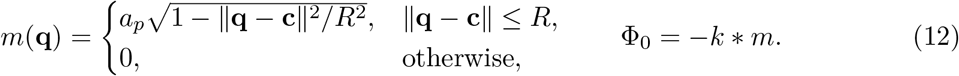

Its response depth is

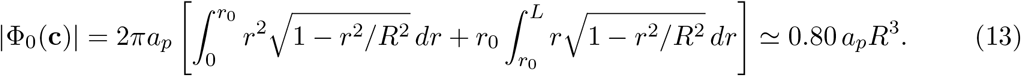

Thus the ratio of the well depth to the pointwise noise standard deviation is approximately 1.13(*a*_*p*_*/σ*)*R*.

The curvature formula (8) gives

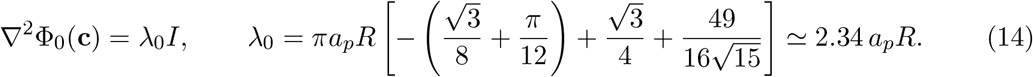

Numerical quadrature of the radial convolution supports the quadratic growth condition (6) of Proposition 1 with

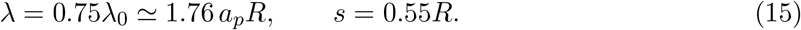

We check these parameters numerically for this idealized continuous profile, since they are not a general bound for experimental particle profiles. For illustration, assume explicitly that

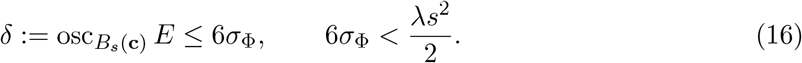

Under these conditions, Proposition 1 gives

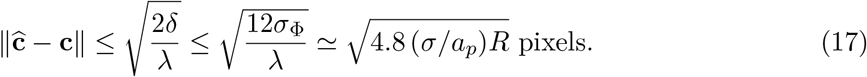

The second condition is equivalent to (*a*_*p*_*/σ*)*R >* 16; for *R* = 82 and *a*_*p*_*/σ* = 1*/*4 it holds with (*a*_*p*_*/σ*)*R* = 20.5, and the bound is approximately 40 pixels, or 0.49*R*, within the radius *s* = 0.55*R*. For this fixed oscillation budget, the relative bound decreases as ((*a*_*p*_*/σ*)*R*)^*−*1*/*2^.

The condition *δ* ≤ 6*σ*_Φ_ is an illustrative oscillation budget, not a probabilistic conclusion from a pointwise noise estimate.

## Footnotes

1 The surrogate function-based centering of (Heimowitz et al., 2021) expands each projection image (extracted picked particle) in a basis of two-dimensional prolate spheroidal wave functions (PSWFs). This expansion must be done separately for each extracted projection image. For this reason, the runtime of such a grid-based picking using the centering method in (Heimowitz et al., 2021) would be long. Therefore, in this paper, we provide a fast alternative with a related local centering criterion of the surrogate function of (Heimowitz et al., 2021) and use it for particle picking.

2 The response can be normalized by a positive scalar (such as *R*^2^) as this will leave all comparisons and extrema unchanged. The quantitative bounds in §3 use the slope-one kernel in (2).

